# Increased α-2,8-sialyltransferase 8B (ST8SIA2) expression in schizophrenia superior temporal gyrus

**DOI:** 10.1101/377770

**Authors:** Toni M. Mueller, Stefani D. Yates, James H. Meador-Woodruff

## Abstract

Reduced polysialylation of neural cell adhesion molecule (NCAM) in schizophrenia has been suggested to contribute to abnormal neuroplasticity and neurodevelopmental features of this illness. The posttranslational addition of sialic acid is mediated by sialyltransferases, and polysialylation (the addition of ≥ 8 α -2,8-linked sialic acid residues) is catalyzed by three enzymes: ST8SIA2 (also called STX), ST8SIA4 (also called PST), and/or ST8SIA3. ST8SIA2 and ST8SIA4 are the primary mediators of NCAM polysialylation. The gene encoding ST8SIA2 maps to schizophrenia risk locus 15q26, and single nucleotide polymorphisms (SNPs) and SNP haplotypes of the ST8SIA2 gene have been associated with schizophrenia in multiple populations. The current study in elderly schizophrenia (N = 16) and comparison (N = 14) subjects measured the protein expression of NCAM, polysialylated-NCAM (PSANCAM), and three poly-α-2,8-sialyltransferases (ST8SIA2, ST8SIA3, and ST8SIA4) in postmortem superior temporal gyrus. Although expression of NCAM, PSA-NCAM, ST8SIA3, and ST8SIA4 were not different in schizophrenia, increased protein levels of ST8SIA2 were identified. It has been reported that ST8SIA2 mutations associated with increased schizophrenia risk impair PSA-NCAM synthesis, suggesting that increased protein expression of ST8SIA2 may represent a compensatory mechanism in the face of impaired enzyme function. This interpretation is further supported by our finding that the relationship between ST8SIA2 enzyme expression and PSA-NCAM levels are different between schizophrenia and comparison subjects. Together these findings suggest a possible neurodevelopmentally-regulated mechanism which could contribute to abnormal synaptic plasticity evident in schizophrenia.

---

Dysregulation of neurotransmission-associated molecules across multiple brain regions, cell types, and neurotransmitter systems has been implicated in the pathophysiology of schizophrenia. We previously hypothesized that deficits of neurotransmission are not only due to abnormal expression of neurotransmission-associated molecules, but may also arise from disrupted posttranslational modifications (PTMs) on substrates involved in key cell signaling processes. Glycosylation is a common PTM that is central to cell signaling, intracellular trafficking, and neuron development. Multiple types of glycosylation and expression levels of glycosylation-associated enzymes are abnormal in schizophrenia.^1–7^

Polysialylation is a specific form of glycosylation and has been previously reported to be abnormal in schizophrenia brain.^1,7^ Polysialylation is the addition of eight or more sialic acid residues (polysialic acid; PSA) to a terminal sialic acid by an α-2,8-linkage, and PSA chain lengths can range from 8 to >100 sialic acid residues attached to a single N-glycan.^8^ In mammals, there are three α-2,8-polysialyltransferases that synthesize PSA chains: ST8SIA2 (also called ST8SiaII, SIAT8B, or STX), ST8SIA3 (ST8SiaIII or SIAT8C), and ST8SIA4 (ST8SiaIV, SIAT8D, or PST).^9–11^ The most well-characterized substrate of this PTM is neural cell adhesion molecule (NCAM), a transmembrane protein that plays essential roles in neurodevelopment, cell migration, synaptic integrity, dendritic morphology, and neuroplasticity.^12,13^ Splice variants of NCAM produce isoforms of 180, 140, and 120kDa (NCAM180, NCAM140, and NCAM120) and non-polysialylated NCAM180 is more sensitive to an extracellular proteolytic cleavage event that produces an additional 105-115kDa isoform (c-NCAM) than NCAM carrying a PSA chain (PSA-NCAM).^11,14,15^ The glycosyltransferases ST8SIA2 and ST8SIA4 primarily mediate the polysialylation of NCAM, though *in vitro* studies suggest that ST8SIA3 may also nominally catalyze glycopeptide polysialylation.^9,10,16,17^ Studies in model systems have found that expression of ST8SIA2 and/or ST8SIA4 can produce the same number of PSA chains on NCAM, and that these enzymes can compensate for one another when mutated forms of either are expressed. The length of PSA chains produced by ST8SIA4 is greater than those produced by ST8SIA2.^9,18,19^

PSA-NCAM expression, but not unmodified NCAM isoform expression, is decreased and c-NCAM—the cleavage product of non-polysialylated NCAM180—is increased in schizophrenia brain.^1,20^ Further, increased c-NCAM expression levels in cerebrospinal fluid (CSF) of patients correlates with ventricular volume increases, suggesting that proteolysis of NCAM may contribute to morphological differences evident in schizophrenia brain.^15^ Studies assessing genetic risk factors of psychiatric illness in multiple geographic populations (Spanish, Chinese, Japanese, Australian, and Canadian) have shown that both the chromosomal region 15q26 and specific haplotypes of the gene encoding ST8SIA2 are associated with schizophrenia and bipolar disorder.^21–25^ Single nucleotide polymorphisms (SNPs) and SNP haplotypes of ST8SIA2 associated with schizophrenia risk result in abnormal ST8SIA2 expression and function, as well as decreased PSA levels *in vitro.*^19,26,27^ While evidence of both upstream regulator dysregulation (genetic mutations of ST8SIA2) and downstream effectors (reduced PSA-NCAM abundance) have been reported in schizophrenia, potential alterations of polysialyltransferase protein levels which could link these findings have not previously been assessed. In this study we measured protein expression of ST8SIA2 and ST8SIA4 in the left superior temporal gyrus (STG; Brodmann area 22) of postmortem schizophrenia brain obtained from elderly subjects in later stages of the disorder. In addition to ST8SIA2 and ST8SIA4, which are known to mediate NCAM polysialylation, we also measured the expression of ST8SIA3 and isoforms and glycoforms of NCAM: PSA-NCAM, NCAM180, NCAM140, and NCAM120.

## Results

### ST8SIA2 expression is increased in schizophrenia STG

We measured the protein expression of the polysialytransferases ST8SIA2-4, as well as isoforms and glycoforms of NCAM (NCAM180, NCAM140, NCAM120, and PSA-NCAM) in the left STG of schizophrenia and comparison subjects. We detected a 19.5% increase in ST8SIA2 protein levels in schizophrenia [t (28) = 2.47, p = 0.019; Figure 1, Table 1]. *Post hoc* testing revealed that ST8SIA2 levels negatively correlate with subject age; however, the difference between diagnostic groups remained significant when covarying for age [F (1,27) = 4.33, p = 0.047]. ST8SIA2 protein levels assessed in rats treated with haloperidol did not differ from vehicle treated animals, suggesting that increased ST8SIA2 expression in schizophrenia is unlikely to be an effect of chronic antipsychotic treatment (Figure 2). Contrary to previous findings in schizophrenia prefrontal cortex and hippocampus,^1,7^ we did not identify any difference in the expression of NCAM isoforms (NCAM180, NCAM140, and NCAM120) or PSA-NCAM in the STG (Table 1). We also did not detect a difference in ST8SIA3 or ST8SIA4 protein levels in schizophrenia (Table 1).

**Figure 1.**
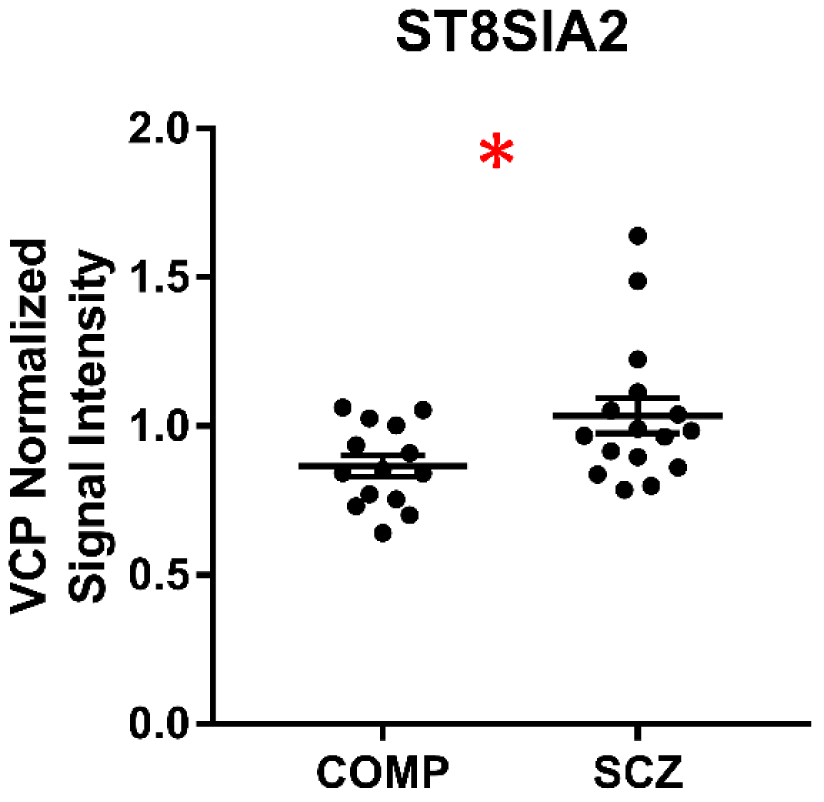
Expression of ST8SIA2 in STG of schizophrenia (SCZ) and comparison (COMP) subjects. Protein expression levels of ST8SIA2 are increased in schizophrenia. Data are presented as mean signal intensity of ST8SIA2 normalized to the signal intensity of intralane VCP for each subject with the mean and S.E.M. of each diagnostic group indicated. *p < 0.05

**Table 1.** Protein Expression.

| Target Protein | Comparison | Schizophrenia | Test statistic (d.f.) | p-value |
| --- | --- | --- | --- | --- |
| ST8SIA2 | 0.866 ± 0.137 | 1.035 ± 0.238 | $t(28) = 2.47$ | 0.019 |
| ST8SIA3 | 0.487 ± 0.088 | 0.510 ± 0.070 | $t(28) = 0.79$ | |
| ST8SIA4 | 0.723 ± 0.142 | 0.756 ± 0.129 | $t(27) = 0.64$ | |
| NCAM (180kDa) | 2.309 ± 0.298 | 2.182 ± 0.375 | $t(27) = 0.29$ | |
| NCAM (140kDa) | 2.195 ± 0.411 | 2.088 ± 0.263 | $t(27) = 0.78$ | |
| NCAM (120kDa) | 3.204 ± 0.513 | 3.164 ± 0.326 | $t(27) = 0.25$ | |
| PSA-NCAM | 1.302 ± 0.155 | 1.317 ± 0.176 | $t(24) = 0.23$ | |

Values represent the mean VCP normalized signal intensity ± the standard deviation (arbitrary units) for each diagnostic group.

**Figure 2.**
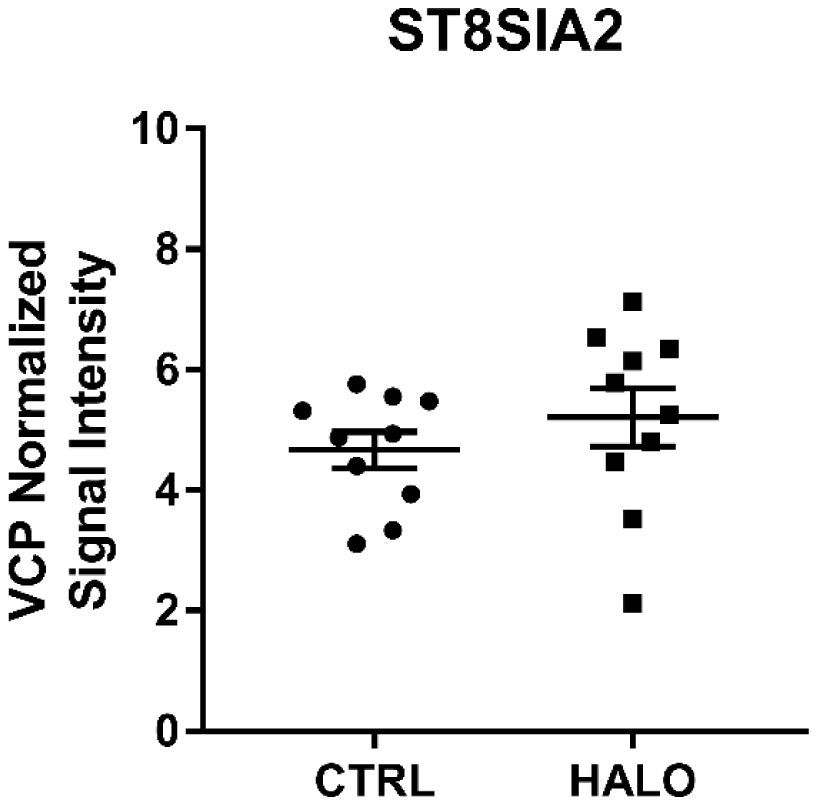
Expression of ST8SIA2 in cortex of rats chronically treated with either haloperidol (HALO) or vehicle (CTRL). There is no difference in the protein expression of ST8SIA2 between HALO or CTRL rats. Data are presented as the mean signal intensity of ST8SIA2 normalized to the signal intensity of intralane VCP with error bars indicating the S.E.M. for each treatment condition.

### The relationship between ST8SIA2 expression and PSA-NCAM levels is altered in schizophrenia STG

For each diagnostic group, linear regression of PSA-NCAM by ST8SIA2 expression was performed and the slope and intercept of best-fit regression lines compared. Neither group demonstrated a significant deviation from slope = 0, however, the slopes of the group regression lines [slope_(SCZ)_ = 0.61 ± 0.37; slope_(COMP)_ = -0.48 ± 0.32) were significantly different [F (1,22) = 4.95; p = 0.037; Figure 3].

**Figure 3.**
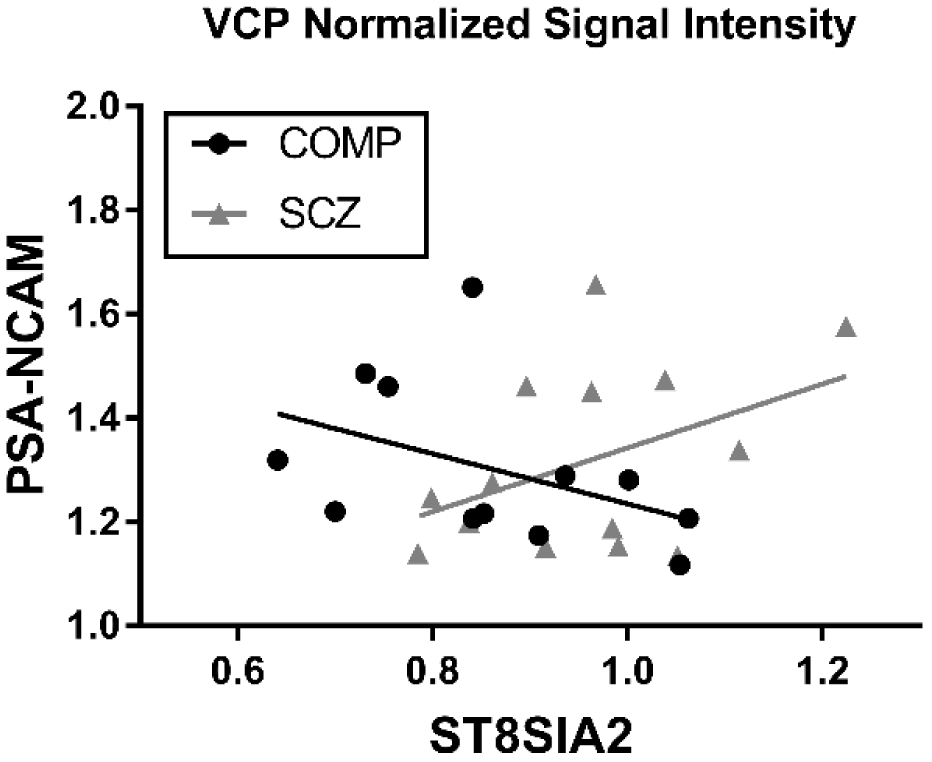
The relationship between ST8SIA2 expression and PSA-NCAM levels is altered in schizophrenia STG. The slopes of best-fit lines of the linear regression of PSANCAM by ST8SIA2 expression for each diagnostic group are significantly different (p = 0.037) and of opposite valence between groups. Data are presented as a scatter plot of the mean signal intensity of target proteins normalized to intralane VCP for each subject.

## Discussion

The current study found increased protein expression of ST8SIA2 and an abnormal relationship between levels of ST8SIA2 and PSA-NCAM in the face of normal NCAM isoform and glycoform protein expression in schizophrenia. The antiadhesive properties of large, flexible, negatively-charged PSA chains on NCAM facilitate structural changes associated with synaptogenesis, axon migration, and neurite outgrowth. PSA-NCAM expression represents an important regulator of spatiotemporal aspects of neuroplasticity during critical neurodevelopmental periods and throughout life, and polysialyltransferase activity is necessary for PSANCAM synthesis.^11–12,13,28^ Prior evidence of reduced NCAM polysialylation without a concurrent reduction in NCAM isoform expression in schizophrenia hippocampus might normally suggest that polysialyltransferase protein levels are reduced. However, the *in vitro* finding that known schizophrenia-associated ST8SIA2 gene mutations result in decreased enzyme function, along with the current finding of increased ST8SIA2 protein levels in the STG, instead suggest that more ST8SIA2 may be expressed to correct for decreased enzyme function in schizophrenia.

Dynamic regulation of PSA-NCAM levels in adult brain is mainly associated with highly neuroplastic brain regions, but NCAM polysialylation also mediates some functions in sensory processing areas. The STG is an important area for auditory processing, and neuroimaging studies show that this region demonstrates cortical asymmetry with volume reductions and reduced regional homogeneity in the left hemisphere in schizophrenia.^29–31^ Consistent with the neuropathology of schizophrenia, an ST8SIA2 deficient mouse model (*St8sia2*^-/-^) demonstrates brain-region specific neuroanatomical alterations and abnormal brain connectivity. *St8sia2*^*-/-*^mice also exhibit working memory defects, decreased social interaction, and reduced prepulse inhibition (PPI) in response to auditory stimuli.^32–34^ Since NCAM polysialylation influences synaptic alterations associated with learning and memory in adults, it is possible that lower levels of PSA-NCAM previously reported may only be evident in in adult brain regions with a very high degree of neuroplasticity and may not be detectible in the STG. Polysialyltransferase-deficient mice exhibit variable patterns of PSA-NCAM expression across multiple brain regions and the resulting increase or decrease in PSA-NCAM level depends not only on which brain region is evaluated but also which enzyme (ST8SIA2 or ST8SIA4) has been knocked out.^34^ Interestingly, NCAM has been shown to require core fucosylation (the addition of α-1,6-linked fucose) prior to PSA chain synthesis by ST8SIA2.^17^ We have recently found that the expression of fucosyltransferase 8 (FUT8), the sole α-1,6-fucosyltransferase expressed in mammals, as well as total levels of core fucose are reduced in these same schizophrenia subjects.^4^ If polysialylation of NCAM is impaired by deficient core fucosylation along with diminished ST8SIA2 function in schizophrenia, as genetic studies suggest, PSA-NCAM levels in schizophrenia brain may not be as sensitive to fluctuations in glycosyltransferase protein expression.

Independent of other possible glycosylation abnormalities, a defect in PSA synthesis by catalytically-impaired ST8SIA2 in schizophrenia could result in chronic upregulation of the dysfunctional ST8SIA2 enzyme while homeostatic levels of PSA-NCAM are maintained in these subjects via the activity of the normally-expressed ST8SIA4 enzyme. Together with evidence of reduced ST8SIA2 efficacy when schizophrenia-associated gene mutations are expressed *in vitro*, the increased level of ST8SIA2 protein we identified likely represents a compensatory upregulation of the enzyme to maintain sufficient homeostatic levels of PSA-NCAM in adult STG. This is further supported by our observation that in elderly schizophrenia subjects, there is a positive relationship between ST8SIA2 and PSA-NCAM expression while the opposite is observed in non-psychiatrically ill subjects. Given that ST8SIA4 can compensate for defects in ST8SIA2 expression and is more functionally efficient than ST8SIA2,^9,19,34^ these findings may also indicate that sufficient levels of PSA-NCAM are more easily maintained in adult non-psychiatrically ill subjects with fully functional ST8SIA2 and ST8SIA4 enzymes.

To reconcile our current report with prior observations, it should be noted that earlier reports of decreased PSA-NCAM in schizophrenia hippocampus and amygdala were based on immunohistochemical staining.^1^ Our study measured all isoforms and glycoforms of NCAM using Western blot analysis, thus it is possible that the difference in methodology could contribute to the disparate results between studies. However, given the cell-type specificity inherent to N-glycan synthesis and processing, we find it more likely that brain-region specific defects in schizophrenia are the main factor underlying this apparent discrepancy.

Limitations of the current study include the relatively small sample size and the unbalanced distribution of subject sex between diagnostic groups. Although replication of these findings with a larger subject set is certainly warranted, the sample size of the current tranche of subjects is sufficiently powered to detect medium effect sizes.

Prior reports of decreased PSA-NCAM expression in the hippocampus and schizophrenia-associated gene mutations producing defects in ST8SIA2 polysialyltransferase function led us to investigate protein levels of polysialyltransferase enzymes, NCAM isoforms, and PSA-NCAM in schizophrenia STG. Despite the limitations, our finding of increased ST8SIA2 protein expression adds to a growing body of literature implicating abnormal PTM processing—specifically, protein glycosylation—in the pathophysiology of the disorder and suggests a possible link between upstream gene mutations and downstream neuroplastic abnormalities in the disorder. Future studies to assess polysialylation in additional brain regions and specific cell types over the course of illness progression may provide insight into regulatory mechanisms that contribute to neurodevelopmental features of schizophrenia.

## Methods

### Subjects and Tissue Acquisition

Postmortem brain tissue was obtained from the Icahn School of Medicine at Mount Sinai NIH Brain and Tissue Repository, and detailed information regarding assessment is available at http://icahn.mssm.edu/research/labs/neuropathologyHYPERLINK-and-brain-banking/neuropathology-evaluation. Brains were evaluated micro- and macroscopically using CERAD guidelines and had no neuropathology or evidence of neurodegenerative disorders, including Alzheimer’s disease, at assessment.^35,36^ Subject exclusion criteria included death not due to natural causes, previous history of drug/alcohol abuse, or coma longer than six hours prior to death.^37^ Subjects used in this study included elderly patients diagnosed with schizophrenia using DSMIII-R criteria, and individuals with no documented history of psychiatric illness obtained from the same collection served as a comparison subjects (Tables 2 and 3). Whole brains were collected at time of autopsy and the full thickness of grey matter from the left STG (Brodmann area 22) was dissected, blocked into 1 cm cubes, nitrogen-pulverized, and stored at -80°C until use. Samples were reconstituted in 1x isotonic extraction buffer (ER0100, Sigma-Aldrich, St Louis, MO) and homogenized on ice in a glass-teflon homogenizer. Homogenate was transferred into a nitrogen cavitation vessel (Parr Instrument Company, Moline, IL) and pressurized for 8 min at 450 psi. Nitrogen-cavitated homogenates were collected through the vessel outlet during decompression.^37^ Protein concentrations using a BCA protein assay kit (Thermo Scientific, Waltham, MA) were obtained for each of the homogenates prior to storage at -80°C.

**Table 2.** Subject Summary.

|  | Comparison | Schizophrenia |
| --- | --- | --- |
| *n* | 14 | 16 |
| Age | 79.4 ± 9.3 | 75.8 ± 11.9 |
| Sex | 4M/10F | 11M/5F |
| PMI (hours) | 10.0 ± 7.3 | 11.4 ± 4.4 |
| Tissue pH | 6.3 ± 0.2 | 6.4 ± 0.3 |
| On/Off Rx | 0/14 | 5/11 |

Abbreviations: Male (M), Female (F), Postmortem interval (PMI), Antipsychotic medication (Rx). Values are expressed as means ± standard deviation. Off Rx indicates patients that had not received antipsychotic medications for 6 weeks or more at time of death.

**Table 3.** Subject Demographics.

| Comparison |  |  |  | Schizophrenia |  |  |  |  |
| --- | --- | --- | --- | --- | --- | --- | --- | --- |
| Sex | Age | Tissue pH | PMI (hrs) | Sex | Age | Tissue pH | PMI (hrs) | Rx |
| F | 73 | 6.30 | 3.4 | F | 53 | 5.88 | 6.3 | on |
| F | 79 | 6.60 | 6.8 | F | 70 | 6.51 | 13.2 | on |
| F | 80 | 6.20 | 4.8 | F | 77 | 6.01 | 9.7 | on |
| F | 80 | 6.32 | 3.8 | F | 83 | 7.10 | 20.4 | off |
| F | 82 | 6.10 | 5.7 | F | 89 | 6.20 | 9.6 | on |
| F | 82 | 6.31 | 24.0 | M | 57 | 6.40 | 15.7 | on |
| F | 84 | 6.21 | 18.5 | M | 66 | 6.50 | 12.1 | on |
| F | 85 | 6.30 | 4.3 | M | 66 | 6.69 | 8.4 | on |
| F | 88 | 6.40 | 5.1 | M | 68 | 6.27 | 8.9 | on |
| F | 89 | 6.72 | 19.0 | M | 73 | 6.30 | 11.7 | off |
| M | 58 | 6.17 | 12.3 | M | 76 | 6.70 | 16.6 | off |
| M | 69 | 6.67 | 7.4 | M | 82 | 6.74 | 11.4 | on |
| M | 70 | 5.86 | 20.5 | M | 84 | 6.71 | 17.7 | off |
| M | 93 | 6.41 | 4.2 | M | 85 | 6.30 | 5.3 | on |
|  |  |  |  | M | 86 | 6.20 | 5.3 | on |
|  |  |  |  | M | 97 | 6.50 | 9.3 | on |

Abbreviations: Female (F), Male (M), Postmortem interval (PMI), Antipsychotic medication (Rx). Off Rx indicates schizophrenia patients that had not received antipsychotic medications for 6 weeks or more at time of death.

### Antipsychotic-treated Rats

Male Sprague-Dawley rats were housed in pairs and treated with chronic administration of either 28.5mg/kg haloperidol decanoate (HALO; N = 10) or vehicle (CTRL; N = 10) delivered once every 3 weeks over 9 months via intramuscular injection, for a total of 12 injections. This method of drug delivery, length of treatment, and dose have been previously described.^3,38,39^Animals were euthanized by rapid decapitation following CO_2_ administration; samples of frontal cortex were dissected on ice then snap frozen and stored at -80°C until homogenization. Homogenates were prepared in 320 mM sucrose in 5 mM Tris-HCL, pH 7.5, with protease and phosphatase inhibitor tablets (Complete Mini, EDTA-free and PhosSTOP, Roche Diagnostics, Indianapolis, IN), and protein concentration determined by BCA Assay (Thermo Scientific) prior to storage at -80°C. The Institutional Animal Care and Use Committee of the University of Alabama at Birmingham approved all procedures using these animals.

### Western Blot Analysis

For both human and rodent samples, homogenates were thawed on ice then prepared with 6x reducing buffer (4.5% sodium dodecyl sulfate (SDS), 0.02% bromophenol blue, 15% β-mercaptoethanol, and 36% glycerol in 170舁mM Tris-HCl, pH 6.8) to a final 1x buffer concentration and heated at 70°C for 10 min. Samples (10 μg/lane) were loaded in duplicate into NuPAGE Bis-Tris 4-12% gradient gels (Life Technologies, Carlsbad, CA) and subjected to SDS-PAGE. Proteins were transferred to nitrocellulose membranes using semi-dry transblotters (Bio-Rad, Hercules, CA). Membranes were blocked and probed using conditions optimized to the linear range of detection for each antibody (Table 4). Membranes were washed with Tris-buffered saline with 0.1% Tween-20 (TBST) and probed with IR-dye labeled secondary antibody in the same diluent as the primary antibody. Membranes were again washed with TBST, rinsed with sterile water, then imaged with a LiCor Odyssey scanner (LiCor, Lincoln, NE). Image Studio Lite Version 4.0.21 (LiCor) was used to measure the signal intensity of each protein band with the median right-left background signal intensity (3 pixels wide) subtracted. Each target was normalized to the intralane signal intensity of valosin-containing protein (VCP), a ubiquitously expressed protein we have previously reported to be unchanged in these same subjects.^37^ Expression levels were similarly assessed in antipsychotic and vehicle treated rats for protein measures that were found altered in schizophrenia.

**Table 4.** Antibody Conditions.

| Target Protein | Host Species | Dilution | Buffer | Incubation | Cat# | Vendor |
| --- | --- | --- | --- | --- | --- | --- |
| ST8SIA2 | rabbit | 1 : 1 000 | 5% BSA | ON 4°C | ab91411 | Abcam |
| ST8SIA3 | rabbit | 1 : 500 | 5% Milk | ON 4°C | 18004-1-AP | Proteintech |
| ST8SIA4 | mouse | 1 : 500 | 50% OBB | ON 4°C | SAB1402975 | Sigma-Aldrich |
| NCAM1 | rabbit | 1 : 500 | 50% OBB | ON 4°C | 14255-1-AP | Proteintech |
| PSA-NCAM | mouse | 1 : 500 | 5% Milk | 72 hr 4°C | MAB5324 | EMD Millipore |
| VCP | mouse | 1:1 000 | 50% OBB | 1 hr RT | ab11433 | Abcam |

Antibodies were obtained from Proteintech (Chicago, IL), EMD Millipore (Billerica, MA), or Abcam (Cambridge, MA). Antibody diluents were adjusted to the indicated concentration with Tris-buffered saline supplemented with 0.1% Tween-20. Abbreviations: bovine serum albumin (BSA); nonfat dry milk (Milk); Odyssey Blocking Buffer (OBB; #927-40100; LiCor, Lincoln, NE); overnight (ON): room temperature (RT).

### Data Analysis

The investigator who executed experimental protocols was blind to subject diagnosis until study completion, and the investigator who performed statistical analyses was blind to diagnosis during data collection. Sample sizes were determined *a priori* using a power calculation (β = 0.2, π = 0.8) based on prior assessments of glycosylation enzyme protein expression in our lab.^4^ Data analysis was performed using Prism 6.07 (GraphPad Software Inc., La Jolla, CA) and STATISTICA 7.1 (StatSoft Inc., Tulsa, OK). Protein expression levels were calculated as the mean of the VCP-normalized target signal intensity. Data were assessed for outliers using the ROUT method (robust regression and outlier removal) with Q = 1%,^40^ then assessed for normal distribution using the Shapiro-Wilk normality test. Data that were not found normally distributed were log-transformed. Normal distributions of raw and transformed data were verified and between group differences were assessed by two-way unpaired Student’s t-tests. *Post hoc* determination of potential associations between enzyme protein levels and PSA-NCAM expression were assessed for each diagnostic group using linear regression and Pearson correlation analyses. For significantly different protein measures, assessments were performed including simple regression analysis between protein expression and subject age, tissue pH, and postmortem interval (PMI); variables demonstrating any significant associations were further assessed by analysis of covariance (ANCOVA) with the correlated measure as covariate. Tests of significant dependent measures grouped by medication status and sex were performed using the non-parametric Kruskal-Wallis test due to small group sizes and no significant associations were identified. Additionally, significantly different dependent measures were assessed in haloperidol treated (N =10) or vehicle control (N = 10) Sprague-Dawley rats by two-way unpaired Student’s ttest following confirmation of normal distribution. For all statistical tests, α = 0.05.

## Acknowledgements

The authors would like to acknowledge the technical expertise of Ms. Anita Pinner and express our sincere gratitude for her assistance in the generation of data. We also gratefully acknowledge the helpful feedback and insight provided by Ms. P Mueller, Dr. W Mueller, and Mr. MR Baker.

## Competing Interests

The authors have no competing interests to disclose.

## Contributions

TMM, SDY, and JMW designed the study. SDY executed experiment protocols. TMM performed data calculations, managed literature searches, performed statistical analyses, and authored the first draft of the manuscript. All authors contributed to and have approved the final manuscript.

## Funding

Funding for this research has been provided by the University of Alabama at Birmingham Department of Psychiatry and Behavioral Neurobiology and the NIH-NIMH under award number R01MH53327.

